# Estimating sleep parameters using an accelerometer without sleep diary

**DOI:** 10.1101/257972

**Authors:** V.T. van Hees, S. Sabia, S.E. Jones, A.R. Wood, K.N. Anderson, M. Kivimäki, T.M. Frayling, A. I. Pack, M Bucan, M.I. Trenell, Diego R. Mazzotti, P. R. Gehrman, B. A. Singh-Manoux, M. N. Weedon

## Abstract

Wrist worn raw-data accelerometers are used increasingly in large scale population research. We examined whether sleep parameters can be estimated from these data in the absence of sleep diaries. Our heuristic algorithm uses the variance in estimated z-axis angle and makes basic assumptions about sleep interruptions. Detected sleep period time window (SPT-window), was compared against sleep diary in 3752 participants (range=60-82years) and polysomnography in sleep clinic patients (N=28) and in healthy good sleepers (N=22). The SPT-window derived from the algorithm was 10.9 and 2.9 minutes longer compared with sleep diary in men and women, respectively. Mean C-statistic to detect the SPT-window compared to polysomnography was 0.86 and 0.83 in clinic-based and healthy sleepers, respectively. We demonstrated the accuracy of our algorithm to detect the SPT-window. The value of this algorithm lies in studies such as UK Biobank where a sleep diary was not used.

---

Wrist-worn raw-data accelerometers are increasingly used for the assessment of physical activity in large population studies such as the Whitehall II study or mega-cohorts such as UK Biobank ^1–3^. The decision to use raw-data accelerometers is motivated by the improved comparability of output across different sensor brands ^4,5^ and better control over all steps in data processing^6^. Accelerometers are commonly worn for 24 hours per day, thus providing information over the day and night; making them potentially valuable for sleep research.

A major challenge in accelerometer-based sleep measurement is to derive sleep parameters without additional information from sleep diaries ^1,3,7^ Standard methods for sleep detection based on conventional accelerometers (actigraphy) involves asking the participant to record their time in bed, sleep onset, and waking up time^8–10^. In a previous paper we developed a method to detect sleep guided by sleep diary records ^11^. However, the increasing use of accelerometry in studies worldwide without sleep diaries necessitates the development of novel methods to derive indicators of sleep behaviour, in the absence of sleep diary records. A crucial step is the detection of the sleep period time window (SPT-window), which is the time window starting at sleep onset and ending when waking up after the last sleep episode of the night. Once the SPT-window can be detected without a diary, our previously published method can be used to detect sleep episodes within this window ^11^. Polysomnography (PSG) is considered the gold-standard measure of sleep parameters, making it an ideal methodology to validate sleep detection methods using an accelerometer algorithm. Additionally, experiments in daily life can be used to establish concurrent validity with sleep diary.

We aim to develop and evaluate a heuristic algorithm for the detection of the SPT-window from raw data accelerometers unaided by a sleep diary and to compare sleep parameters (waking up, sleep onset time and SPT-window duration) with sleep diary records assessed in the daily life of a large cohort of older adults, and with PSG data collected in a sleep clinic and a group of healthy good sleepers.

## Methods

### Study population

In order to assess the validity of our algorithm in different settings and against both data from sleep diary and polysomnography, data are drawn from three different study populations described below.

The Whitehall II cohort study^12^: full details on data collection were previously described ^11^. Briefly, accelerometer measurement was added to the study at the 2012/2013 wave of data collection for participants seen at the central London clinic and for those living in the South-Eastern regions of England who underwent a clinical evaluation at home ^2^ Of the 4879 participants to whom the accelerometer was proposed in the Whitehall II Study, 388 did not consent and 210 had contraindications (allergies to plastic or metal, travelling abroad the following week). Of the remaining 4281 participants who wore the accelerometer, 4204 (98.2%) had valid accelerometer data (a readable data file). Among them, sleep diary data were missing for 80 participants and 29 additional participants did not meet criteria for accelerometer wear time (at least one night defined as noon-noon with >16h of wear time). Of the remaining 4095 participants (a total of 27,966 nights) 342 did not have complete demographic data (age, BMI and sex). Therefore, the main assessment of discrepancies between the accelerometer and the sleep diary was undertaken in 3752 participants (76.9% of those invited) with a total of 25,645 nights ^11^. The resulting participants (75.2% men) were on average 69.1 (standard deviation (SD) = 5.6) years old and had a mean body mass index (BMI) of 26.4 (SD = 4.2) kg/m^2^.

Sleep clinic patients: these data come from 28 adult patients who were scheduled for a one-night polysomnography (PSG) assessment at the Freeman Hospital, Newcastle upon Tyne, UK, as part of their routine clinical assessment and were subsequently invited to participate in the study ^11^. All 28 patients recruited for the polysomnography study (11 female) had complete accelerometer data for the left wrist and 27 had complete data for the right wrist and were aged between 21 and 72 years (mean±sd: 45±15 years). Diagnosed sleep disorders included: hypersomnia (N=2), insomnia (N=2), REM behaviour disorder (N=3), sleep apnoea (N=5), narcolepsy (N=1), sleep apnoea (N=4), parasomnia (N=1), restless leg syndrome (N=5), and sleep paralysis (N=1), and nocturnia (N=1). Three patients had more than one sleep disorder.

Healthy good sleepers: these data come from 22 adults who underwent a one-night PSG assessment at the University of Pennsylvania Center for Sleep. Twenty-two participants recruited for the polysomnography study (68% female) had complete accelerometer data for the non-dominant wrist and were aged between 18 and 35 years (mean±sd: 22.8±4.5 years).

### Ethics Statement

In all three studies participants were provided with instructions and an information sheet about the study and were given time to ask questions prior to providing written informed consent. The studies were approved by the University College London ethics committee (85/0938) and the NRES Committee North East Sunderland ethics committee (12/NE/0406), and University of Pennsylvania ethics committee (819591) respectively. All experiments were performed in accordance with relevant guidelines and regulations.

### Data availability

Whitehall II data, protocols, and other metadata are available to the scientific community. Please refer to the Whitehall II data sharing policy at https://www.ucl.ac.uk/whitehallII/data-sharing. Raw data from the polysomnography study has been made open access available in anonymized format on zenodo.org^13^. Data from the University of Pennsylvania are available through the National Institute of Mental Health data archive.

### Instrumentation

Participants in the Whitehall II Study were asked to wear a tri-axial accelerometer (GENEActiv, Activinsights Ltd, Kimbolton, UK) on their non-dominant wrist for nine (24-h) consecutive days. They were asked to complete a simple sleep diary every morning which consisted of two questions: ‘what time did you first fall asleep last night?’ and ‘what time did you wake up today (eyes open, ready to get up)?’ The accelerometer was configured to collect data at 85.70 Hz with a ±8g dynamic range. A more complete description of the accelerometer protocol can be found in our earlier publication ^2^.

In the second and third study, polysomnography (Embletta®, Denver) was performed using a standard procedure, including video recording, a sleep electroencephalogram (leads C4-A1 and C3-A2), bilateral eye movements, submental EMG, and bilateral anterior tibialis EMG to record leg movements during sleep. Respiratory movements were detected with chest and abdominal bands measuring inductance, airflow was detected with nasal cannulae measuring pressure, and oxygen saturation of arterial blood was measured. Airflow limitation and changes in respiratory movement were used to detect increased upper-airway resistance. All respiratory events and sleep stages were scored according to standard criteria so that EEG determined total sleep time could be measured ^9^. Participants in the second study (PSG in sleep clinic) were asked to wear the same brand of accelerometer as in the first study (GENEActiv, Activinsights Ltd, Kimbolton, UK) on both wrists throughout the one-night polysomnography assessment. Here, the accelerometer was also configured to record at 85.70 Hz. Accelerometer data were collected on both wrist to assess the role of sensor location on classification performance, unfortunately no information on handedness was recorded. Participants in the third study (PSG in healthy good sleepers) were asked to wear an accelerometer of the brand Axivity (Axivity Ltd, Hoults Yard, UK) on the non-dominant wrist throughout the one-night polysomnography assessment. Here, the accelerometer was configured to record at 100 Hz.

### Accelerometer data preparation

A previously published method was used to minimize sensor calibration error ^14^ and to detect and impute accelerometer non-wear periods ^2,15^. Arm angle was estimated as follows: 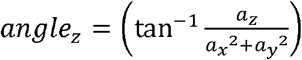, where *a*_*x*_, *a*_*y*_, and *a*_*z*_ are the median values of the three orthogonally positioned raw acceleration sensors in gravitational (*g*) units (1*g* = 1000 mg) derived based on a rolling five second time window. Here, the z-axis corresponds to the axis positioned perpendicular to the skin surface (dorsal-ventral direction when the wrist is in the anatomical position). Next, estimated arm angles were averaged per 5 second epoch and used as input for our algorithms for detecting sleep period time (SPT-window) and sleep episodes.

### Heuristic algorithm to detect the SPT-window

There are several challenges in the development of an algorithm to detect the SPT-window: absence of hard data labels to train a classifier under daily life conditions (not in a clinic), consideration of daily life behaviour, e.g. how to handle sleep scattered across the full 24-hour day and ensure that the algorithm is not over fitted to a specific population or accelerometer brand. Thus an algorithm was developed by visually inspecting twenty random accelerometer multi-day recordings from different studies and accelerometer brands (ten from the Whitehall II Study as reported in this paper and ten from UK Biobank study ^1^) while iteratively enhancing the algorithm to best detect the visible data segment of no movement without using or looking at sleep diary data.

The resulting heuristic algorithm, which we will refer to as Heuristic algorithm looking at Distribution of Change in Z-Angle (HDCZA), applied per participant is illustrated in Figure 1 and works as follows. *Step 1-2:* Calculate the z-angle per 5 seconds. *Steps 3-5:* Calculate a 5-minute rolling median of the absolute differences between successive 5 second averages of the z-angle. These first five steps make the algorithm invariant to the potentially unstandardized orientation of the accelerometer relative to the wrist and aggregate it as the rolling variance over time. *Step 6-7:* Calculate the 10^th^ percentile from the output of step 5 over an individual day (noon-noon), and multiply by 15. This is used as a critical individual night derived threshold to distinguish periods of time involving many and few posture changes. Detect the observation blocks for which the output from step 5 was below the critical threshold, and keep the ones lasting longer than 30 minutes. *Step 8:* Evaluate the length of the time gaps between the observation blocks identified by step 7, if the duration is less than 60 minutes then count these gaps towards the identified blocks. *Step 9:* The longest block in the day (noon-noon) will be the main SPT-window, defined as the time elapsed between sleep onset (start of the block) and waking time (end of the block). These last four steps reflect assumptions from us as researcher about the nature of sleep.

**Figure 1:**
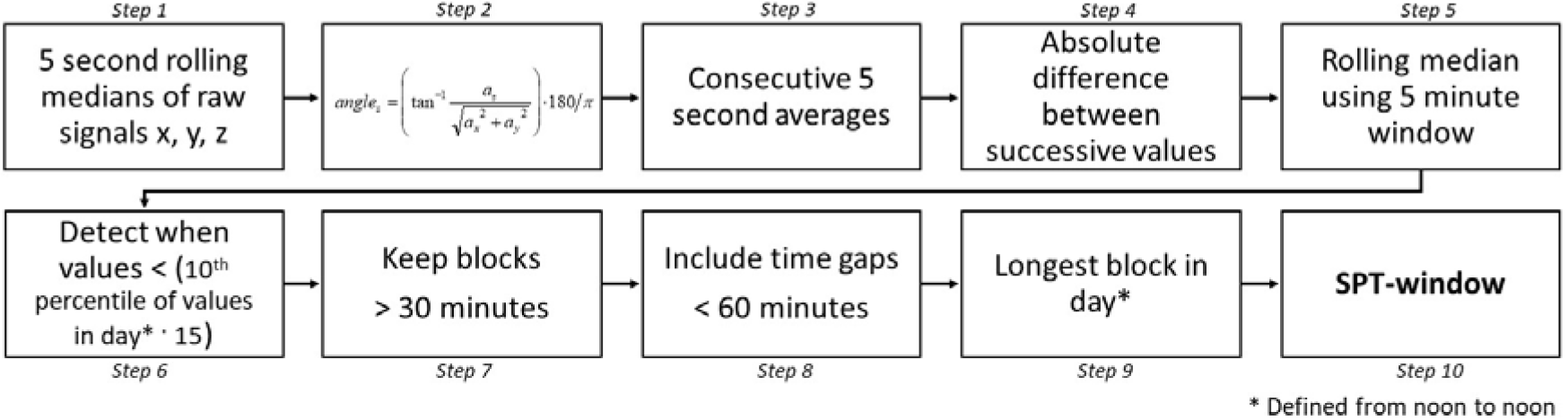
Steps of the heuristic algorithm HDCZA for SPT-window detection.

Our motivation for the design of the algorithm is as follows. By visually inspecting the angle-z values over a day some individuals seemed inactive or sleeping throughout the day with minimal variation in angle, while other individuals had more distinct inactive (night time) and active (daytime) periods. These differences presumably reflect the degree of sedentary lifestyle and amount of sleep in a day. Using a percentile as part of the threshold calculation allows the threshold to account for between-individual differences in z-angle distribution. The factor 15 in step 6 of the algorithm was derived iteratively using visual inspection of the classification. The 30-minute time period is motivated by the assumption that people are typically not in bed for less than 30 minutes for their nocturnal time in bed, as opposed to daytime napping, and the 60-minute time period is motivated by the assumption that sleep separated by awake periods greater than 60 minutes ought to be treated as two distinct sleep episodes to avoid adding early evening naps or afternoon naps to the SPT-window. A sensitivity analysis on HDCZA parameter settings and their influence on algorithm performance across the datasets can be found in Supplementary material 3.

### Second algorithm for reference

When comparing our algorithm to the sleep diary we also considered a second, but more naïve heuristic algorithm, which we will refer to as L5±6. The algorithm is based on the raw signal metric Euclidian Norm (vector magnitude) Minus One with negative values rounded to zero (ENMO), which in formula corresponds to 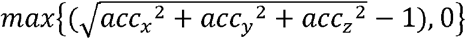, with acc_x_, acc_y_, and acc_z_ referring to the three orthogonal acceleration axes pointing in the lateral, distal, and ventral directions, respectively ^15^. Metric ENMO has previously been demonstrated to be correlated with magnitude of acceleration as well as human energy expenditure in the present generation of wearable acceleration sensors^15^. L5±6 takes the 12 hour window centred around L5 (least active five hours in the day based on metric ENMO) and then searches within this window for sustained inactivity periods which were previously described ^11^. In short, sustained inactivity periods are calculated as the absence of change in arm elevation angle (same angle-z as used above) larger than 5 degrees for more than 5 minutes ^11^. Next, the SPT-window is defined from the start of the first to the end of the last occurrence of a sustained period of inactivity in the 12-hour window.

### Sleep episodes within the SPT-window

Sleep episodes were defined as the sustained periods of inactivity within the SPT-window, as defined in the previous section ^11^. From this, the number of sleep episodes within each SPT-window detected (HDCZA, L5±6) was calculated as well as sleep efficiency within the SPT-window calculated as the percentage of time asleep within the SPT-window ^11^.

## Statistical analysis

### Comparison with sleep diary

The SPT-window derived from both the HDCZA and L5±6 were compared separately with sleep diary records with a multi-level regression to account for the variation in availability of night time data and to include both night and person level predictors. For SPT-window duration (difference between sleep onset and waking time), sleep onset and waking time, the difference between diary and accelerometer-based detection was used as the dependent variable, while population demographics (sex, age, BMI), season (winter or summer) and weekend versus weekday were used as predictors. Here, we used function lme from R package nlme. Further, correlation coefficients and mean absolute error (MAE) between sleep onset, waking time, and SPT-window duration were calculated. Additionally, the c-statistic, also known as the Area Under the Curve (ROC), was calculated from the epoch-level binary classifications of SPT-window <1> or not <0> by diary and the HDCZA and L5±6, first calculated per day and then aggregated as average per participant. Additionally, to investigate whether more wakefulness time within the SPT-window corresponds to a larger HDCZA-sleep diary difference in SPT-window duration we calculated the amount of wakefulness categorised as [0-1), [1-2), [2-3), [3-4), and at least 4 hours, and compared this with the difference in SPT-window duration between sleep diary and the HDCZA. The notation [a-b) is used to denote an interval that is inclusive of ‘a’ but exclusive of ‘b’.

### Evaluation with polysomnography

The recording time of PSG is typically constrained to the time in bed window, which means that our heuristic algorithm (HDCZA) may not detect sufficient data corresponding to time out of bed to derive its critical threshold and accurately detect the SPT-window. We addressed this concern by adding simulated wakefulness data to the beginning and ending of the accelerometer and PSG recording. The PSG and accelerometer data were expanded with 90 minutes of simulated data at the beginning and ending that would not trigger the SPT-window detection: simply the class wakefulness for PSG, and a sine wave with amplitude 40 degrees and period 15 minutes complemented with random numbers (mean=0, standard deviation=10) for accelerometer-based angle-z. Note that the specific shape of the simulated values is not critical as long as it does not trigger the detection of sleep and the 10^th^ percentile of all the data (step 6 of HDCZA) reflects real and not simulated data. The addition of simulated data is needed because the heuristic detection algorithm effectively searches for the beginning and end of a large time period without body movement, if the full PSG represents sleep then the algorithm would not be able to detect such a transition in movement level. Additionally, the algorithm’s threshold that scales with the variance in the data was constrained to a range corresponding to the 2.5^th^ and 97.5^th^ percentile of the distribution of the threshold value observed in a sample of daily life accelerometer recordings, 0.13 and 0.50, respectively. This was done because the in-clinic PSG does not provide a full 24-hour cycle of body movement to derive this threshold. In the PSG evaluation we did not evaluate L5±6, because it requires more than 12 hours of (non-simulated) data, which most PSG recordings do not offer. After sleep classification with HDCZA and before running the comparison between HDCZA and PSG, 60 minutes of simulated data were removed at the beginning and end.

The following performance metrics for SPT-window detection were used: difference in onset, waking time, and duration, accuracy, c-statistic, t-test, and mean absolute error (MAE). Performance estimates accuracy and c-statistic were derived from both the data, as well as from the data expanded with wakefulness time to simulate performance estimates in a 24 hour recording. Sleep classification within the SPT-window was evaluated as difference in duration (t-test) and as the percentage of time spent in sleep stages REM, and non-REM stages 1, 2, and 3 (N1, N2, and N3) correctly classified by the algorithm as part of SPT-window. Sleep efficiency within the SPT-window by PSG and algorithm was compared via t-test and MAE. A *P*-value of < .005 was considered significant^16^. Further, method agreement was evaluated with modified Bland-Altman plots^17^ with PSG criterion values on the horizontal axis.

### Code availability

Both SPT-window detection algorithms are implemented and available in open source R package GGIR version 1.5-21 (https://cran.r-project.org/web/packages/GGIR/)^18^, see the software’s documentation on input arguments ‘loglocation’ and ‘def.noc.sleep’ for further details on the use of L5±6 and HDCZA. The R code used for our comparisons with sleep diary can be found at: https://github.com/wadpac/whitehall-acc-spt-detection-eval. The R code used for our comparisons with polysomnography can be found at: https://github.com/wadpac/psg-ncl-acc-spt-detection-eval, with the code used for the Newcastle data in the master branch of the repository and its adaptation for the differently formatted Pennsylvanian data in the psg-penn branch.

## Results

### Comparison between accelerometer results and that from sleep diary

Demographic characteristics of the three study cohorts are described in Table 1. The probability density distribution for the difference between sleep parameter estimates from algorithm and sleep diary is more symmetrical around zero compared with the L5±6 approach, see Figure 2. The heuristic algorithm HDCZA estimates sleep onset on average 12.5 and 7.5 minutes earlier than that reported in the sleep diaries by men and women, respectively, 3.9 minutes per ten years of age relative to mean age, and 3.0 minutes for a weekend day, see Table 2. Difference between sleep diary estimates and HDCZA estimates in waking time and SPT window duration were associated with sex, age, and BMI, see Table 2. The L5±6 method estimates sleep onset on average 86.4 and 78.5 minutes earlier than that reported in the sleep diary for men and women, respectively. Difference between sleep diary and L5±6 estimates of SPT-window, sleep onset, and waking time were associated with sex and BMI, but inconsistently with weekday, see Table 2. The Pearson’s correlation coefficients and c-statistics between accelerometer derived sleep parameters, and sleep diary, are higher for HDCZA compared with L5±6, see Table 3. The combined MAE from onset and waking time was 34.8 and 75.6 minutes for HDCZA and L5±6, respectively.

For nights with [0-1), [1-2), [2-3), [3-4), and at least 4 hours of accumulated wakefulness an average difference in SPT-window duration between sleep diary records and our heuristic algorithm (HDCZA) was observed as 27, 3, −58, −154, and −236 minutes corresponding to 57.9, 32.1, 7.5, 1.6, and 0.7% of 25,645 recorded nights, respectively. Here, the last two categories, corresponding to at least 3 hours of accumulated wakefulness, reflect 8.5% of the participants.

**Table 1:** Participant characteristics used for the analyses

| **Study** | **Daily life (diary)** | **PSG sleep clinic** | **PSG healthy good sleepers** |
| --- | --- | --- | --- |
| N | 3752 | 28 | 22 |
| Age (mean ± standard deviation in years) | 69.1 ± 5.6 | 44.9 ± 14.9 | 22.8 ± 4.5 |
| Sex | 2822 males, 930 females | 17 males and 11 females | 7 males and 15 females |
| SPT-window duration (mean ± standard deviation) | 7.7 ± 1.2 hours | 8.4 ± 1.6 hours | 6.7 ± 0.9 hours |
| Sleep onset time (mean in hh:mm ± standard deviation) | 23:48 ± 71 minutes | 22:32 ± 69 minutes | 23:24 ± 54 minutes |
| Waking time (mean in hh:mm ± standard deviation) | 7:28 ± 72 minutes | 06:58 ± 76 minutes | 06:09 ± 32 minutes |

**Figure 2:**
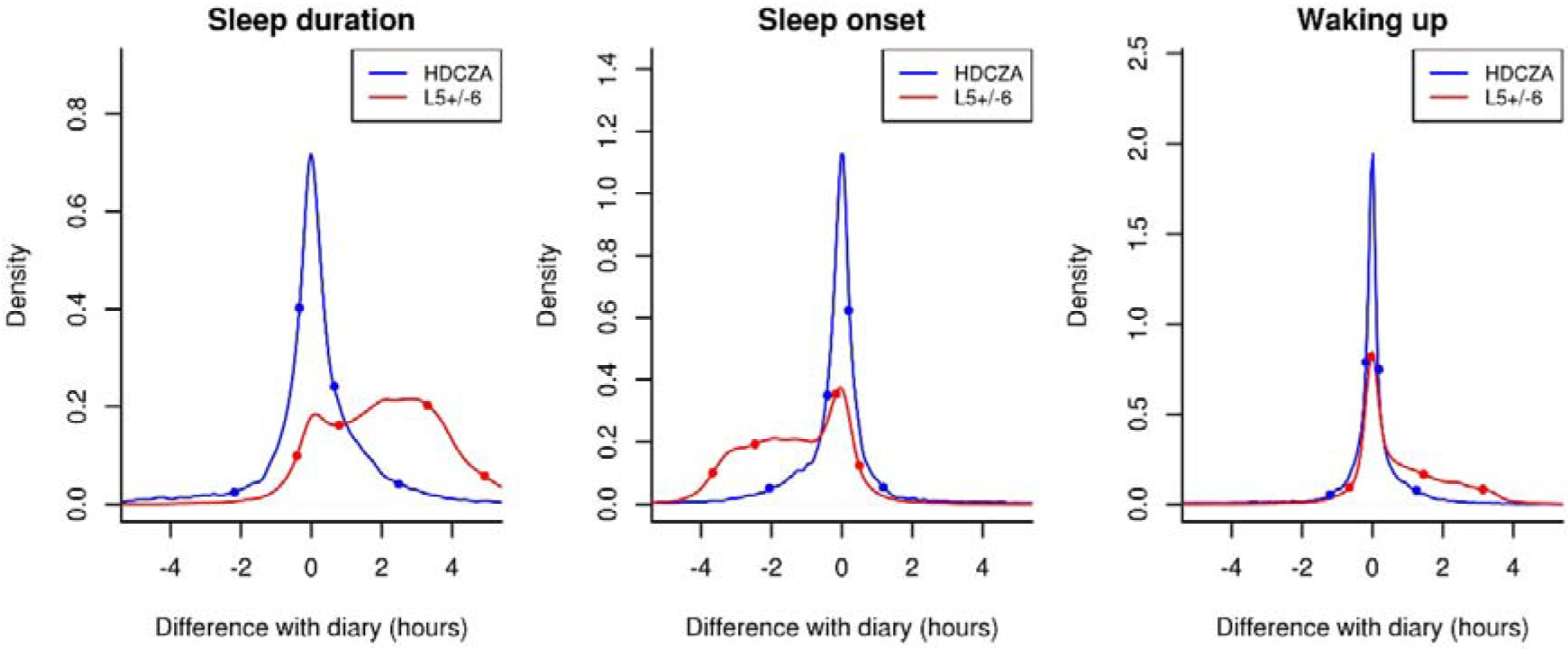
Probability density distributions for accelerometer-based estimates of sleep duration, sleep onset, and waking up time using dots to indicate the 5^th^, 25^th^, 75^th^ and 95^th^ percentile.

**Table 2:** Sleep parameter differences (minutes) between estimates from sleep diary and two accelerometer-based methods (N=25,645 nights, N=3752 individuals)

| Sleep parameters | HDCZA |  |  | L5±6 |  |  |
| --- | --- | --- | --- | --- | --- | --- |
| Method | Sleep onset time | Waking time | SPT-window duration | Sleep onset time | Waking time | SPT-window duration |
| **Y-intercept (SE)** | -12.5 (0.9) ** | -1.6 (0.8) P=0.04 | 10.9 (1.1) ** | -86.5 (1.0) ** | 45.9 (0.9) ** | 131.7 (1.2) ** |
| **Betas (SE)** |  |  |  |  |  |  |
| Women | 5.0 (1.1) ** | -3.0 (0.9) * | -8.0 (1.3) ** | 8.0 (1.4) ** | -8.6 (1.1) ** | -16.2 (1.6) ** |
| Ten years of age † | 3.9 (0.8) ** | -2.9 (0.7) ** | -6.8 (1.0) ** | 0.3 (1.0) P=0.78 | 0.2 (0.8) P=0.83 | -0.2 (1.2) P=0.89 |
| Five BMI index points ‡ | 1.0 (0.5) P=0.06 | -1.5 (0.5) * | -2.5 (0.7) ** | -3.2 (0.7) ** | 1.8 (0.6) * | 4.8 (0.8) ** |
| Weekend | 3.0 (1.0) * | 2.0 (0.9) P=0.02 | -1.0 (1.2) P=0.41 | 6.4 (1.3) ** | -0.3 (1.0) P=0.77 | -6.3 (1.4) ** |
| Winter | 1.0 (0.9) P=0.27 | -1.2 (0.8) P=0.12 | -2.2 (1.1) P=0.05 | -1.0 (1.2) P=0.39 | 0.6 (1.0) P=0.51 | 1.7 (1.3) P=0.2 |
| Within individual residual SD | 24.7 | 21.3 | 30.9 | 18 | 13 | 20.6 |
| Between individual residual SD | 66.1 | 56.9 | 82.3 | 88.9 | 74.9 | 101.8 |
| AIC | 81175 | 73538 | 92433 | 94053 | 84956 | 100978 |

[Degrees of freedom=25,645; † relative to mean age of 69.1 years; ‡ relative to mean BMI of 26.4 kg / m<sup>2</sup>; SE: Standard Error; SD: Standard Deviation; AIC = Akaike information coefficient, \* P < .005, \*\* P < .0005]

**Table 3:** Correlation, mean absolute error, and concordance between sleep diary and accelerometer estimates (N=3,752)

| Parameter | Metric | HDCZA |  |  | L5±6 |  |  |
| --- | --- | --- | --- | --- | --- | --- | --- |
|  |  | Value | t; DF | P | Value | t; DF | P |
| sleep onset time | Correlation in timing | 0.78 (95% CI: 0.77 - 0.79) | 76; 3750 | ** | 0.66 (95% CI: 0.64 - 0.68) | 54; 3750 | ** |
|  | MAE (min) | 39.9 |  |  | 93.3 |  |  |
| waking time | Correlation in timing | 0.81 (95% CI: 0.8 - 0.82) | 84; 3750 | ** | 0.68 (95% CI: 0.66 - 0.7) | 57; 3750 | ** |
|  | MAE (min) | 29.9 |  |  | 58.4 |  |  |
| SPT-window | Correlation in duration | 0.52 (95% CI: 0.5 - 0.55) | 38; 3750 | ** | 0.26 (95% CI: 0.23 - 0.29) | 16; 3750 | ** |
|  | MAE (min) | 40 |  |  | 128.4 |  |  |
|  | c-statistic | 0.95 (IQR: 0.94 - 0.98) | - | - | 0.92 (IQR: 0.90 - 0.94) † | - | - |

[DF: Degrees of freedom; MAE: mean absolute error; min: minutes; \* P < 0.005; \*\* P < 0.0005; † -0.03 difference (95% CI for difference: -0.031; -0.029), t=-44, DF=3751, P < .0005]

### Comparison between accelerometer results and that from polysomnography

In the PSG study in sleep clinic patients, on average 9.4 (standard deviation 1.6) hours of matching data from PSG and accelerometer were retrieved per participant, with no difference in recording duration between left and right wrist (P = 0.75). Sleep onset time, waking time, SPT-window duration, and sleep duration within the SPT-window derived from the HDCZA algorithm differed all non-significantly from polysomnography and MAE ranged from 31 minutes for sleep onset to 71 minutes for SPT-window duration, see Table 4. The combined MAE from onset and waking time was 38.9 and 36.7 minutes for the left and right wrist, respectively. SPT-window duration was estimated for the left wrist within 2 hours for the majority of individuals (75 %) but deviated by more than 2 hours in seven individuals, six of which had a sleep disorder, as shown in Figure 3 (right wrist: 81%, five, and four, respectively). On average, the accuracy and C-statistic for SPT-window classification were 87% and 0.86 in the PSG recording window, and 94% and 0.94 when expanded with simulated wakefulness as an estimate of 24 hour performance, see Table 4. Further, the average sensitivity to detect sleep as part of the SPT-window was above 91% in both wrists, see Table 4. Results for the PSG study carried out in healthy good sleepers indicated better overall performance as shown in Table 5 and Figure 4. The classifications of the HDCZA algorithm in comparison with the PSG sleep stage classification for all participants are provided in the Supplementary material 1 and 2 to this manuscript.

**Figure 3:**
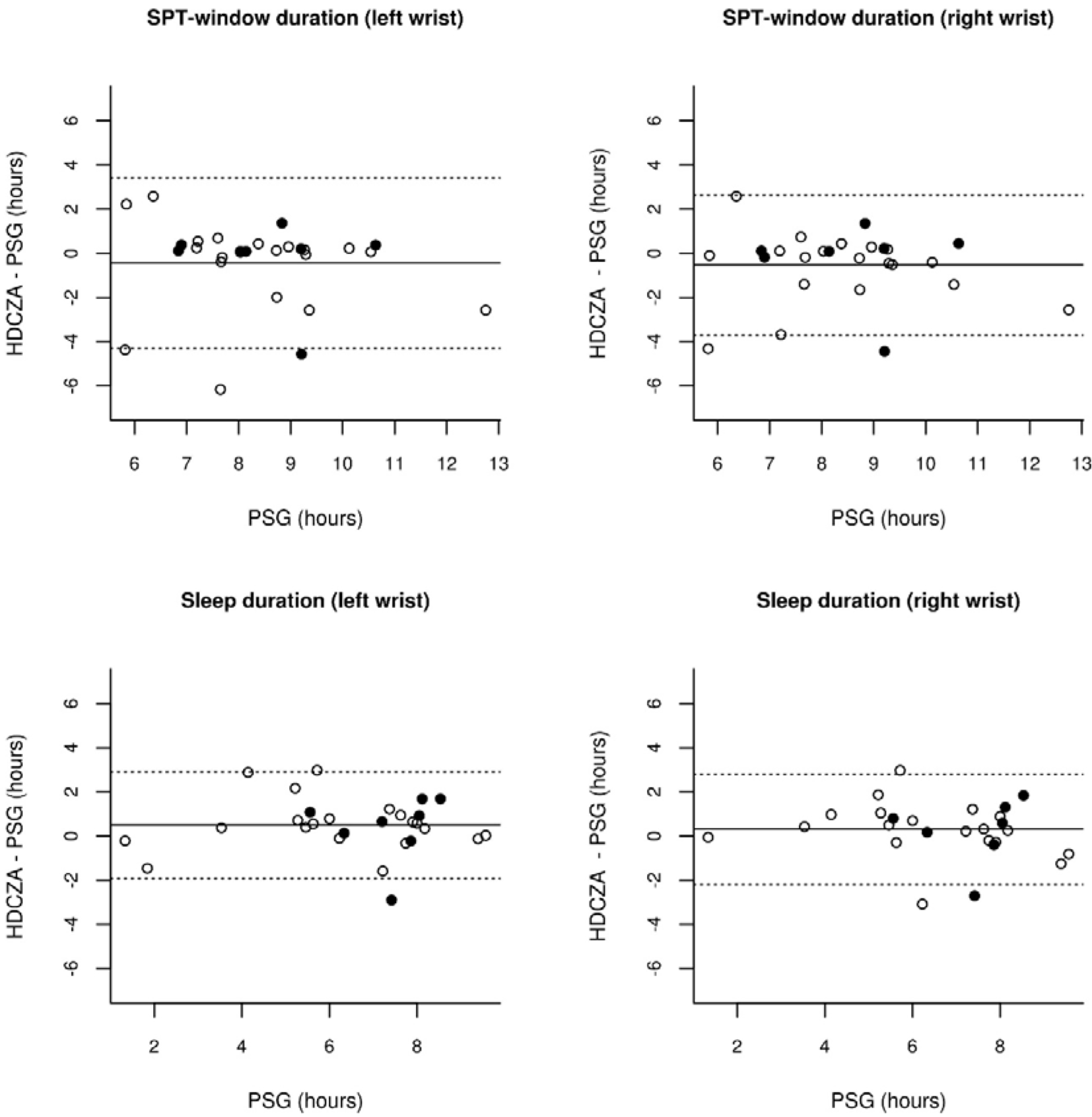
Modified Bland-Altman plots with 95% limits of agreement (LoA) for SPT-window duration and sleep duration relative to polysomnography (PSG) in sleep clinic patients, with dashed lines indicating LoA and straight line indicating the mean. Open bullets reflect individuals with a sleep disorder, while closed bullets reflect normal sleepers.

**Figure 4:**
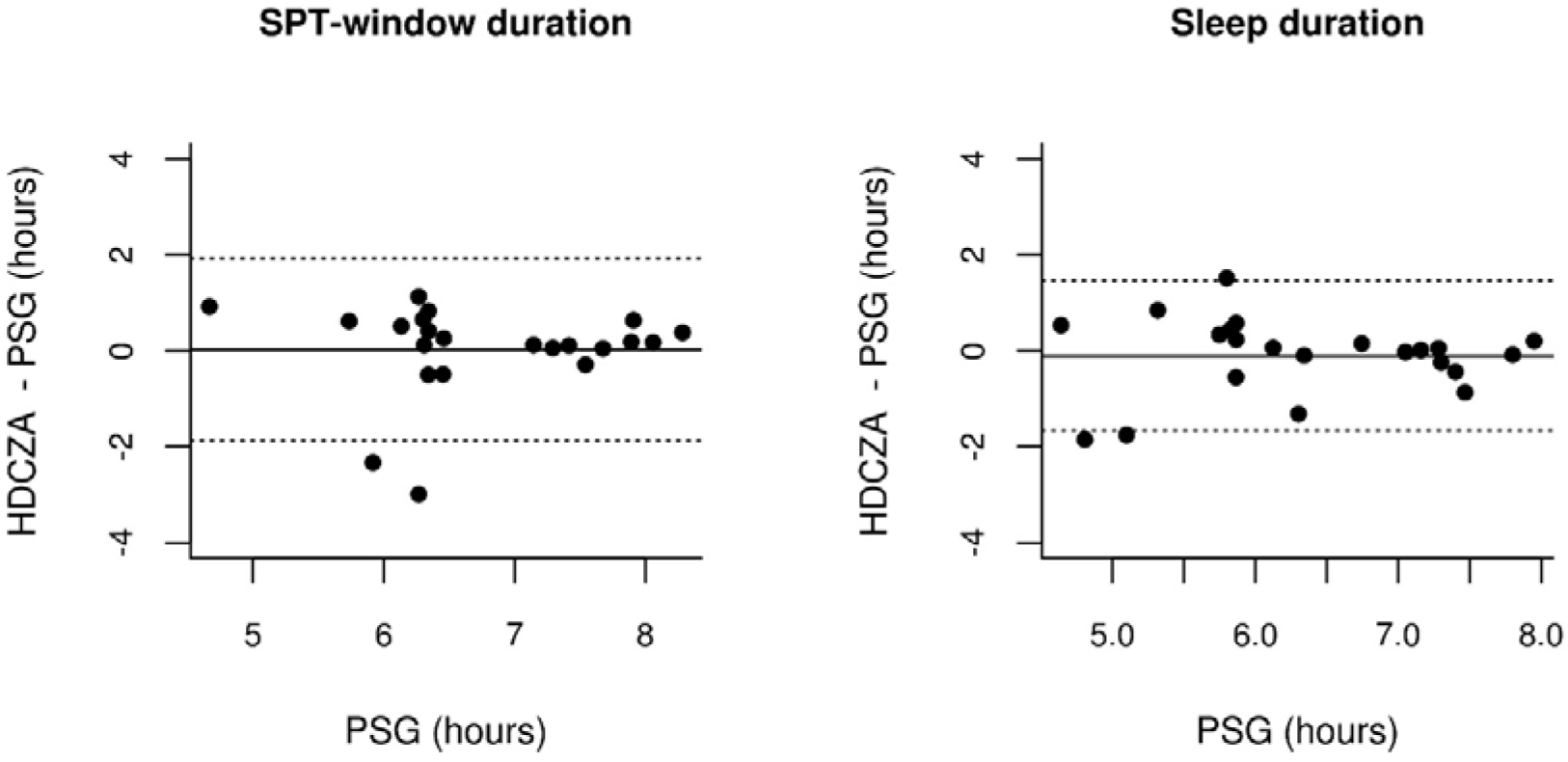
Modified Bland-Altman plots with 95% limits of agreement (LoA) for SPT-window duration and sleep duration relative to polysomnography (PSG) in healthy good sleepers, with dashed lines indicating LoA and straight line indicating the mean.

**Table 4:** Comparison algorithm with polysomnography in sleep clinic patients (Newcastle study)

| Parameters | Metric | Left wrist (N=28) |  |  | Right wrist (N=27) |  |  |
| --- | --- | --- | --- | --- | --- | --- | --- |
|  |  | Value | t; DF | P | Value | t; DF | P |
| Sleep onset | Difference (min) | -10 (95% CI: -30; -9) | -1.08; 27 | 0.29 | 0 (95% CI: -27; 27) | 0.02; 26 | 0.98 |
|  | MAE (min) | 30.8 | - | - | 40.2 |  |  |
| Sleep wake | Difference (min) | -37 (95% CI: -75; 1) | -2.00; 27 | 0.06 | -31 (95% CI: -57; -6) | -2.54; 26 | 0.02 |
|  | MAE (min) | 47.1 | - | - | 33.2 |  |  |
| SPT-window | Difference in duration (min) | -27 (95% CI: -73; 19) | -1.21; 27 | 0.23 | -32 (95% CI: -71; 6) | -1.72; 26 | 0.10 |
|  | MAE (min) | 70.9 | - | - | 63.5 | - | - |
|  | c-statistic | 0.86 (IQR: 0.81-0.98) | - | - | 0.87 (IQR: 0.81-0.95) | - | - |
|  | c-statistic 24 hour† | 0.93 (IQR: 0.94-0.99) | - | - | 0.94 (IQR: 0.94-0.99) | - | - |
|  | Accuracy (%) | 87 (IQR: 81-98) | - | - | 88 (IQR: 84-97) | - | - |
|  | Accuracy 24 hour† (%) | 94 (IQR: 92-99) | - | - | 94 (IQR: 93-99) | - | - |
| Sleep within SPT | Difference in duration (min) | 30 (95% CI: 1; 58) | 2.11; 27 | 0.04 | 18 (95% CI: -12; 48) | 1.24; 26 | 0.23 |
|  | Sensitivity (%) | 92 (IQR: 97-100) | - | - | 91 (IQR: 98-100) | - | - |
| Sleep efficiency within SPT | Difference (percent point) | 8.7 (95% CI: 3.63 – 13.82) | 3.51; 27 | * | 9.4 (95% CI: 3.76 – 15.06) | 3.42; 26 | * |
|  | MAE (percent point) | 10.1 | - | - | 10.6 | - | - |

[\* P < .005; MAE: mean absolute error; min: minutes; SPT-window: Sleep period time window; CI: Confidence Interval; DF: degrees of freedom; t: t-statistic; IQR: Inter quartile range; † recording expanded with simulated data of wakefulness to resemble 24 hours]

**Table 5:**
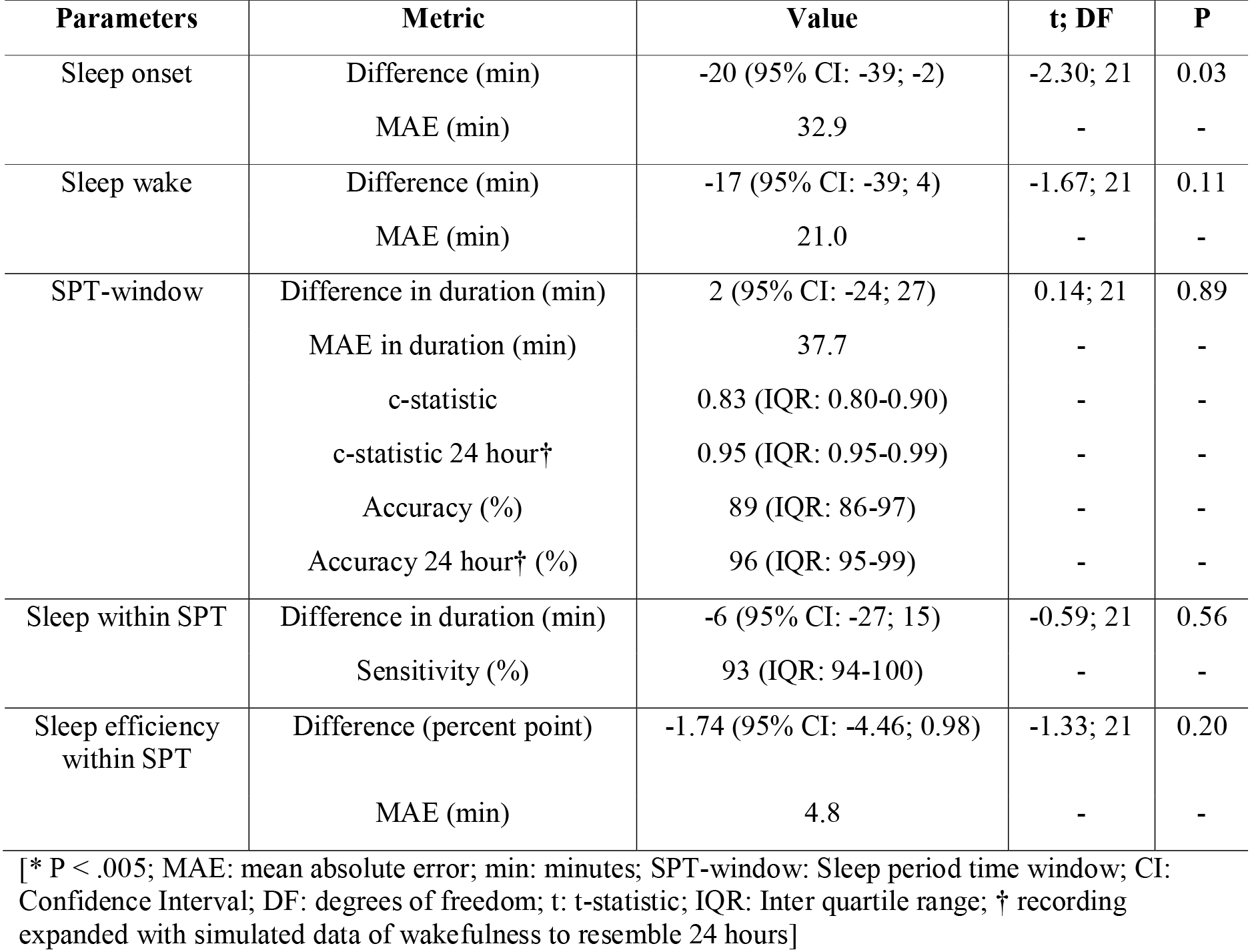
Comparison algorithm with polysomnography in healthy good sleepers (N=22, Pennsylvania)

## Discussion

In this paper we present a heuristic algorithm, referred to as HDCZA, for detecting Sleep Period Time-window (SPT-window) from accelerometer data in the absence of a sleep diary. Raw data accelerometers are increasingly used in population research, and the value of this algorithm lies in studies such as the UK Biobank where a sleep diary was not used ^1^. Although the focus of our analysis is sleep, the present findings are equally valuable for physical activity research as it will help to split the observation period between night sleep and daytime inactivity.

In our comparison with sleep diary records in a large cohort of older adults (60-82 years) a small systematic difference was found in sleep duration and sleep onset time, difference that varies slightly as a function of sex, age, and BMI. Here, the average difference and the Akaike Information Coefficients indicated that the algorithm is better than our naïve reference method L5±6. Furthermore, the C-statistic was on average 95% for HDCZA. We acknowledge that the sleep diary cannot be considered a gold standard criterion method, but it is reassuring to see that differences between algorithm and sleep diary in a large cohort of elderly individuals are on average within a quarter of an hour.

An important limitation of the sleep diary study data is that no information is available on daytime sleep or daytime inactivity behaviour to help better understand the misclassifications in SPT-window by our algorithm. To facilitate such research future methodological studies are warranted to consider implementing daytime sleep diaries, and possibly additional sensor technologies such as wearable cameras^19^, RFID proximity sensors^20^ or additional wearable movement sensors to better capture a lying posture^21,22^. In addition, impact of handedness on the estimates could not be assessed.

When compared against polysomnography in 28 sleep clinic patients, accuracy and C-statistic values indicate good agreement on an epoch by epoch level. Estimated SPT-window duration by HDCZA deviated by more than 2 hours from PSG in seven individuals (six of which has a sleep disorder) as shown in figure 3. Inspection of the PSG results indicated that poor classification typically occurs in patients with absence of deep sleep or who have long periods of wakefulness (> 1 hour) in the middle of the night, e.g. pages 10 and 26 in the Supplementary material, respectively (see Supplement 1). However, the interpretation of the results was complicated in case of SPT-window split into several periods separated by long waking periods. For example, one particular individual had a short sleep episode at the beginning of the PSG recording followed by several hours of wakefulness, see page 9 of the Supplementary material (Supplement 1), indicating a possible ambiguity in the correct definition of the SPT-window by both PSG and HDCZA.

To investigate the extent to which the larger differences in individuals with long periods of wakefulness observed in the PSG study occur in the general population we went back to the free-living data from the first study. In the free-living data, more wakefulness during the night corresponded to larger differences between sleep diary and algorithm derived SPT-window duration, indicating that more wakefulness time is indeed a challenge in a daily life recording setting. However, it was reassuring to see that only a small fraction (2.4%) of all the nights scattered across 8.5% of the participants were affected by one hour or more. In line with this observation the tails in the distribution of differences with sleep diary (Figure 2) may be explained by wakefulness during the night or sleep episodes being scattered over the day. The problem then is that the SPT- window lacks a clear construct definition. Another possible explanation for the tails in the distribution includes the subjective nature of sleep diary as well.

Differences and mean absolute error were better in the evaluation with healthy good sleepers (Pennsylvania), indicating that SPT-window detection is a challenge in those with sleep disorders. The expansion of PSG data with daytime wakefulness to simulate algorithm performance in a full day has to our knowledge not been done before. We think this can help the comparison and interpretation of the c-statistic between the night time only PSG and full day sleep diary studies. A downside of this approach is that it comes with the assumption that daytime is always correctly classified. Therefore, we presented both performance estimates with and without the additional simulated data.

In the absence of a gold standard criterion method that can be applied in a representative part of the population under daily life conditions to train and test a classifier, we consider the heuristic approach the most promising for detecting the SPT-window. The heuristic approach comes with the following advantages: (i) It is not optimized with subjective and therefore potential erroneous sleep diary records, (ii) It avoids potentially overfitting towards a small patient population in a PSG study unrepresentative for the general population, (iii) It does not make assumptions about the timing or duration of the SPT-window, and (iv) It is computationally simple which will facilitate easy replication. The sensitivity analysis on parameter configuration as reported in supplementary material 3 demonstrates that the current configuration provides a relatively good average performance across alternative configurations that is relatively robust against changing study conditions. Improvement in algorithm performance in a specific dataset via optimization of parameter configuration can lead to overfitting, which comes with poor performance in other datasets or a subset of the data.

We found one other study that compared SPT-window extracted from accelerometry (or actigraphy) unaided by sleep diary to facilitate further interpretation of our current findings. Recently, O’Donnell and colleagues also investigated possible approaches to SPT-window detection, currently available as a non-peer reviewed preprint on bioRxiv ^23^. To compare algorithm performance, we replicated their main performance metric: the mean absolute error (MAE) in sleep onset and waking time. Our HDCZA algorithm has a MAE of 34.8 minutes when compared against sleep diary (N=3751), which is comparable with the 33.3, 34.4, and 35.9 minutes reported for the three algorithms investigated by O’Donnell (N=14)^22^. Although the age range is similar between the studies, a substantial difference in sample size and unknown differences in the prevalence of disturbed sleep warrants future standardized comparison between the algorithms. Further, the MAE estimates in our PSG studies are 38.9, 36.7, and 26.9 minutes in the left- and right wrist sleep clinic patient data, and healthy good sleepers, respectively. When we consider the design of our and their approach, we observe a couple of differences: their change-point and random forest approaches were optimized on a trained data set with sleep diary data as criterion, which our approach avoids following aforementioned point (i). Further, O’Donnell’s thresholding approach relies on the assumption that the average SPT-window duration is 8 hours, which our approach also avoids following aforementioned point (iii). Other strengths of our approach are the evaluation with sleep diary in much larger cohort than theirs and we evaluated our approach against PSG in sleep clinic patients arguably a challenging subpopulation to classify sleep in. Neither our nor their approach currently uses the available temperature or light sensor information, in our case because of concerns about measurement bias from environmental conditions. Therefore, future research is needed to explore the potential of temperature and light information to enhance the SPT-window classification.

It should be noted that the historical studies like the one by Cole-Kripke^24^ and later studies ^25,26^ focussed on automatic distinction of sleep and wakefulness aided by the boundaries of time in bed, lights off, or diary records of the SPT-window. These studies then focussed on correct classification of Wake After Sleep Onset (WASO), Total Sleep Time (TST), and Sleep Efficiency. Overall these sleep estimates based on algorithms aided by sleep diary show better agreement with PSG estimates than algorithms not aided by a sleep diary. However, these studies represent a different measurement construct and methodological challenge than discussed in the present work and can therefore not be used as a reference point. To give the reader an idea of how much better the MAE is when a sleep diary is available to aid the detection of the SPT window, we have calculated this from the analysis in our previous publication^11^: the MAE was on average 12 minutes (inter quartile range: 7-15) using the same sleep diary as reference point.

Our algorithm does not facilitate the detection of sleep latency. To derive sleep latency, one would need diary records of time in bed or the lights out period. Future research is warranted to investigate how sleep latency, time in bed, and the lights out period may reliably be detected from wearable accelerometer data without asking the participant to record their sleep behaviour using a diary or marker button.

The analysis presented in this paper will facilitate feasible large-scale population research on sleep and physical activity. In addition to the proof of validity as provided in this paper additional support for the credibility of the algorithm was found in our separate study (non-peer reviewed preprint on bioRxiv) identifying genome wide associations with sleep parameters derived from our algorithm in UK Biobank, replicating signals previously associated with self-reported sleep duration and chronotype ^27–34^ Our algorithm can be applied to data from the three most widely used accelerometer brands: Actigraph, Axivity, and GENEActiv, and is available as part of open source R package GGIR (https://cran.r-project.org/web/packages/GGIR/).

## Acknowledgements

This work was made possible thanks to the following grants: NIH grants HL-094307 (AIP), and; MRC grant MR/P012167/1. We would like to thank Dr. Sarah Charman, Dr. Paul Innerd, Matthew Goodman and Sara McHugh-Grant for their contributions to the collection of PSG data.

## Author contributions

V.T.v.H. ‘contributed to’ conception and design of the work, data analysis and interpretation, article drafting, and critical revision of the article. S.S. and K.N.A. ‘contributed to’ data collection, and critical revision of the article. S.E.J., A.R.W., M.K., T.F., D.R.M., P.G., M.B., M.I.T., B.A.S.M., M.N.W, A.I.P ‘contributed to’ critical revision of the article.

## Additional information

The author(s) declare no competing interests.

## References

1. Doherty, A. et al. Large Scale Population Assessment of Physical Activity Using Wrist Worn Accelerometers: The UK Biobank Study. PLoS One 12, e0169649 (2017).

2. Sabia, S. et al. Association between questionnaire- and accelerometer-assessed physical activity: the role of sociodemographic factors. Am. J. Epidemiol. 179, 781–90 (2014).

3. da Silva, I. C. et al. Physical activity levels in three Brazilian birth cohorts as assessed with raw triaxial wrist accelerometry. Int. J. Epidemiol. 43, 1959–68 (2014).

4. Rowlands, A. V, Yates, T., Davies, M., Khunti, K. & Edwardson, C. L. Raw Accelerometer Data Analysis with GGIR R-package: Does Accelerometer Brand Matter? Med. Sci. Sports Exerc. 48, 1935–41 (2016).

5. Rowlands, A. V et al. Accelerometer-assessed Physical Activity in Epidemiology: Are Monitors Equivalent? Med. Sci. Sports Exerc. 50, 257–265 (2018).

6. van Hees, V. T. et al. Challenges and Opportunities for Harmonizing Research Methodology: Raw Accelerometry. Methods Inf. Med. 55, 525–532 (2016).

7. Anderson, K. N. et al. Assessment of sleep and circadian rhythm disorders in the very old: the Newcastle 85+ Cohort Study. Age Ageing 43, 57–63 (2014).

8. Girschik, J., Fritschi, L., Heyworth, J. & Waters, F. Validation of self-reported sleep against actigraphy. J. Epidemiol. 22, 462–468 (2012).

9. Lockley, S. W., Skene, D. J. & Arendt, J. Comparison between subjective and actigraphic measurement of sleep and sleep rhythms. J. Sleep Res. 8, 175–183 (1999).

10. Littner, M. et al. Practice parameters for the role of actigraphy in the study of sleep and circadian rhythms: an update for 2002. Sleep 26, 337–341 (2003).

11. van Hees, V. T. et al. A Novel, Open Access Method to Assess Sleep Duration Using a Wrist-Worn Accelerometer. PLoS One 10, e0142533 (2015).

12. Marmot, M. G. et al. Health inequalities among British civil servants: the Whitehall II study. Lancet (London, England) 337, 1387–93 (1991).

13. van Hees, V. T., Charman, S. & Anderson, K. N. Newcastle polysomnography and accelerometer data. (2018). doi:10.5281/zenodo.1160410

14. van Hees, V. T. et al. Autocalibration of accelerometer data for free-living physical activity assessment using local gravity and temperature: an evaluation on four continents. J. Appl. Physiol. 117, 738–44 (2014).

15. van Hees, V. T. et al. Separating movement and gravity components in an acceleration signal and implications for the assessment of human daily physical activity. PLoS One 8, e61691 (2013).

16. Benjamin, D. J. et al. Redefine statistical significance. Nat. Hum. Behav. 2, 6–10 (2018).

17. Bland, J. M. & Altman, D. G. Statistical methods for assessing agreement between two methods of clinical measurement. Lancet 1, 307–310 (1986).

18. van Hees, V. et al. GGIR. (2018). doi:10.5281/zenodo.1154149

19. Doherty, A. R. et al. Using wearable cameras to categorise type and context of accelerometer-identified episodes of physical activity. Int. J. Behav. Nutr. Phys. Act. 10, 22 (2013).

20. Shinmoto Torres, R. L., Visvanathan, R., Abbott, D., Hill, K. D. & Ranasinghe, D. C. A battery-less and wireless wearable sensor system for identifying bed and chair exits in a pilot trial in hospitalized older people. PLoS One 12, e0185670 (2017).

21. Bussmann, J. B. J., Veltink, P. H., Koelma, F., van Lummel, R. C. & Stam, H. J. Ambulatory monitoring of mobility-related activities: the initial phase of the development of an activity monitor. Eur. J Phys Med Rehabil 5, 2–7 (1995).

22. Gloeckl, R. et al. Validation of an activity monitor during sleep in patients with chronic respiratory disorders. Respir. Med. 109, 286–8 (2015).

23. O’Donnell, J. et al. Automated detection of sleep-boundary times using wrist-worn accelerometry. (2017). doi:https://doi.org/10.1101/225516

24. Cole, R. J., Kripke, D. F., Gruen, W., Mullaney, D. J. & Gillin, J. C. Automatic sleep/wake identification from wrist activity. Sleep 15, 461–9 (1992).

25. Jean-Louis, G., Kripke, D. F., Mason, W. J., Elliott, J. A. & Youngstedt, S. D. Sleep estimation from wrist movement quantified by different actigraphic modalities. J. Neurosci. Methods 105, 185–91 (2001).

26. Blackwell, T. et al. Comparison of sleep parameters from actigraphy and polysomnography in older women: the SOF study. Sleep 31, 283–91 (2008).

27. Lane, J. M. et al. Genome-wide association analyses of sleep disturbance traits identify new loci and highlight shared genetics with neuropsychiatric and metabolic traits. Nat. Genet. 49, 274–281 (2017).

28. Cade, B. E. et al. Common variants in DRD2 are associated with sleep duration: the CARe consortium. Hum. Mol. Genet. 25, 167–79 (2016).

29. Byrne, E. M., Gehrman, P. R., Trzaskowski, M., Tiemeier, H. & Pack, A. I. Genetic Correlation Analysis Suggests Association between Increased Self-Reported Sleep Duration in Adults and Schizophrenia and Type 2 Diabetes. Sleep 39, 1853–1857 (2016).

30. Jones, S. E. et al. Genome-Wide Association Analyses in 128,266 Individuals Identifies New Morningness and Sleep Duration Loci. PLoS Genet:. 12, e1006125 (2016).

31. Jones, S. E. et al. Genome-wide association analyses of chronotype in 697,828 individuals provides new insights into circadian rhythms in humans and links to disease. (2018). doi:10.1101/303941

32. Dashti, H. et al. GWAS in 446,118 European adults identifies 78 genetic loci for selfreported habitual sleep duration supported by accelerometer-derived estimates. (2018). doi:10.1101/274977

33. Jones, S. E. et al. Genetic studies of accelerometer-based sleep measures in 85,670 individuals yield new insights into human sleep behaviour. (2018).

34. Lane, J. M. et al. Biological and clinical insights from genetics of insomnia symptoms. (2018).

